# Passive muscle stretching reduces estimates of persistent inward current strength in soleus motor units

**DOI:** 10.1101/2020.07.31.230029

**Authors:** Gabriel S. Trajano, Janet L. Taylor, Lucas B. R. Orssatto, Craig R. McNulty, Anthony J. Blazevich

## Abstract

Prolonged (≥60 s) passive muscle stretching acutely reduces maximal force production at least partly through a suppression of efferent neural drive. The origin of this neural suppression has not been determined, however some evidence suggests that reductions in the amplitude of persistent inward currents (PICs) in the motoneurons may be important. The aim of the present study was to determine whether acute passive (static) muscle stretching affects PIC strength in gastrocnemius medialis (GM) and soleus (SOL) motor units. We calculated the difference in instantaneous discharge rates at recruitment and derecruitment (ΔF) for pairs of motor units in GM and SOL during triangular isometric plantar flexor contractions (20% maximum) both before and immediately after a 5-min control period and immediately after five 1-min passive plantar flexor stretches. After stretching there was a significant reduction in SOL ΔF (−25.6%; 95%CI = -45.1 to -9.1 %, p=0.002) but not GM ΔF. These data suggest passive muscle stretching can reduce the intrinsic excitability, via PICs, of SOL motor units. These findings (1) suggest that PIC strength might be reduced after passive stretching, (2) are consistent with previously-established post-stretch decreases in SOL but not GM EMG amplitudes during contraction, and (3) indicate that reductions in PIC strength could underpin the stretch-induced force loss.

**SUMMARY STATEMENT:** Motoneurons require an amplification mechanism to operate within the firing frequencies observed during normal motor behaviour. Here we present evidence that this amplification mechanism is reduced after passive muscle stretching.

## INTRODUCTION

Prolonged (usually ≥60 s; (Behm et al. 2016; Kay and Blazevich 2012)) passive muscle stretching reduces maximal force production and neural drive to the muscle. For example, passive stretching of the human plantar flexors reduces EMG amplitudes during maximal voluntary isometric contractions and both the reduction and recovery of EMG tends to correlate strongly with the loss and recovery of maximal muscle force (Kay and Blazevich 2009; Trajano, Nosaka, et al. 2013; Pulverenti et al. 2019; Ryan et al. 2014). These changes in EMG are strongly associated with changes in other measures of neurological function, including estimates of voluntary activation and V-wave amplitudes (Trajano, Nosaka, et al. 2013; Ryan et al. 2014), leading to the conclusion that muscle force loss largely results from a decrease in neural drive to the muscle.

The origin of this loss of neural drive is still unknown. However, evidence has been provided that passive muscle stretching might cause prolonged reduction in excitability of the alpha motoneurons (Trajano, Nosaka and Blazevich 2017). This conclusion was based on the observation in humans that passive muscle stretching could reduce the strength of the involuntary plantar flexor contractions that normally persist after cessation of simultaneous tendon vibration and electrical muscle stimulation (Trajano et al. 2014). Tendon vibration can elicit muscular contractions via the Ia reflex loop, and these contractions display characteristics that are consistent with the initiation of persistent inward currents (PICs) in α-motoneurons, including self-sustained motor unit firing (after vibration cessation), joint angle dependence, and a warm-up effect (Trajano et al. 2014). Thus, the reduction in strength of involuntary contractions that persist after tendon vibration provides evidence of a stretch-induced disfacilitation of the α-motoneurons, possibly via a reduction in the strength of PICs.

PICs are depolarizing currents generated by voltage-sensitive sodium and calcium channels predominantly located on the motoneurons’ dendrites (Heckman, Gorassini and Bennett 2004). They amplify and prolong synaptic input producing sustained depolarization of the cell, causing motoneurons to continue to fire without need for additional input (Heckman, Gorassini and Bennett 2004; Lee and Heckman 2000). PICs can also change the input-output gain of motoneurons by ∼5-fold or more, depending on the level of monoaminergic drive to the spinal cord, allowing motoneurons to fire at much higher frequencies than when PICs are not triggered (Lee and Heckman 2000). In fact, the neuromodulatory effect exerted by monoamines (i.e. serotonin and noradrenaline in motoneurons) on PICs is so important for normal motor behavior that maximal activation of excitatory ionotropic inputs would produce less than 40% of the maximal motor output in the absence of neuromodulation (Heckman 1994; Johnson et al. 2017). Thus, PICs are a fundamental component of normal motor output observed in humans, and reductions in PIC amplitudes can significantly affect the muscle’s ability to produce force.

In humans, PIC amplitudes are typically estimated using the paired-motor unit technique (Gorassini et al. 2002; Johnson et al. 2017; Wilson et al. 2015). This method measures the discharge rate of a lower-threshold motor unit (control unit), as a surrogate for the level of excitatory drive, at the time of recruitment and de-recruitment of a higher-threshold unit (test unit). The difference between recruitment and de-recruitment frequencies is referred to as the delta frequency (ΔF). ΔF is considered to be proportional to the PIC amplitude and has been validated by comparison with animal data (Powers, Nardelli and Cope 2008) as well as computer simulations (Powers and Heckman 2015). Hence, if passive muscle stretching reduces the ability to develop PICs in humans, as suggested by previous research, then reductions in ΔF should be observed immediately after the stretching has been completed.

Given the above, the main purpose of the present study was to determine whether passive stretching can reduce ΔFs, i.e. estimates of PIC amplitude, in the human plantar flexors. To do this we measured ΔFs in soleus and gastrocnemius medialis before and after 5 sets of 1 min plantar flexor stretching or a control condition (5-min rest). We hypothesized that ΔFs would be lower after passive stretching, indicating a loss of PIC strength.

## MATERIAL AND METHODS

### Participants

Eighteen recreationally active participants (10 males and 8 females) without any neuromuscular limitations volunteered to participate in the study (32±4 y, 73±19 kg, 174±11 cm). Participants were instructed to avoid vigorous exercise and alcohol consumption for 24 h, and caffeine use for at least 6 h, prior to testing. All participants read and signed the informed consent document, and the Queensland University of Technology Human Research Ethics Committee approved this study (Approval number: 1800000550).

### Study design and overview

All data collection was performed in a single session lasting approximately 1 h and 30 min. Participants were seated upright in the chair of an isokinetic dynamometer (Biodex System 4, Biodex Medical system, Shirley, NY) with the knee fully extended (0°) and ankle at neutral position (0°). They were then instructed to practice six voluntary submaximal isometric plantar flexion contractions (2 × 40%, 2 × 60% and 2 × 80% of perceived maximal effort). After this warm-up, two maximal voluntary contractions (MVC) were performed with a 1-min passive rest interval, and maximum torque was recorded. Then, participants practiced isometric triangular ramped contractions, with the instruction to contract their plantar flexors in order to increase torque from 0 to 20% of MVC in 10 s at a rate of 2% of MVC/s, and then decrease that force linearly at the same rate back to the baseline value. Once the participants were familiarized with these triangular contractions, data were recorded as they performed, in the following order, three triangular contractions interspaced by 25 s immediately before (Control 1) and after (Control 2) a 5-min rest period as well as immediately after 5 sets of 1-min passive plantar flexor stretching interspaced by 15-s intervals (Post-stretch).

### Electromyography (EMG)

Surface EMG was recorded during the contractions using two semi-disposable 32-channel electrode grids with a 10-mm inter-electrode distance (ELSCH032NM6, OTBioelettronica, Torino, Italy) placed over the muscle belly of both soleus (SOL) and gastrocnemius medialis (GM). A strap electrode was dampened and positioned around the ankle joint as a ground electrode. The EMG signals were recorded in monopolar mode and converted to digital data by a 16-bit wireless amplifier (Sessantaquattro, OTBioelettronica, Torino, Italy). EMG signals were amplified (256×), sampled at 2000 Hz and bandpass filtered (10-500 Hz) before being stored for offline analysis.

Surface EMG signals were decomposed into single motor unit discharge events using a convolutive blind source separation algorithm using DEMUSE software (Holobar and Zazula 2007). This algorithm has been extensively validated against the standard intramuscular fine wire methods across a variety of muscles and contractions (Holobar, Minetto and Farina 2014; Negro et al. 2016; Muceli et al. 2015). All decomposed motor units were visually inspected and all erroneous identified motor units and discharge times were excluded. Only motor units with a pulse-to-noise ratio above 30dB were kept for further analysis. After that, discharge events with ISIs below 0.025 s and above 0.4 s were excluded. Subsequently, discharge events were converted into instantaneous discharge rates and fitted into a 5^th^-order polynomial function. This polynomial fit was then used for the paired motor unit analysis. ΔF was calculated as the change in discharge rate of a lower-threshold (control) motor unit from the moment of recruitment to the moment of de-recruitment of a higher-threshold (test) unit (Gorassini et al. 2002) (see Figure 1). ΔFs were calculated for pairs of motor units that fitted the following criteria: 1) rate-rate correlations ≥0.7; 2) test unit recruited at least 0.5 s after control unit; and 3) control unit did not show saturation of discharge rates (discharge rates increased by at least 0.5 pulses per second (pps) after the recruitment of the test unit) (Stephenson and Maluf 2011). ΔFs were calculated for individual test units as the average value obtained when the units were paired with multiple control units. Subsequently, a single ΔF value was obtained per participant per muscle (SOL and GM) by averaging the values obtained for all the units from each muscle. Motor units were tracked throughout the conditions using the decomposition filter and the same motor unit pairs were used to calculate the ΔF value for each contraction within each participant. Peak discharge rates were measured as the highest value in the polynomial function for each motor unit and averaged across all the units obtained for each muscle. In three participants (2 females and 1 male) it was not possible to identify usable motor units.

**Figure 1.**
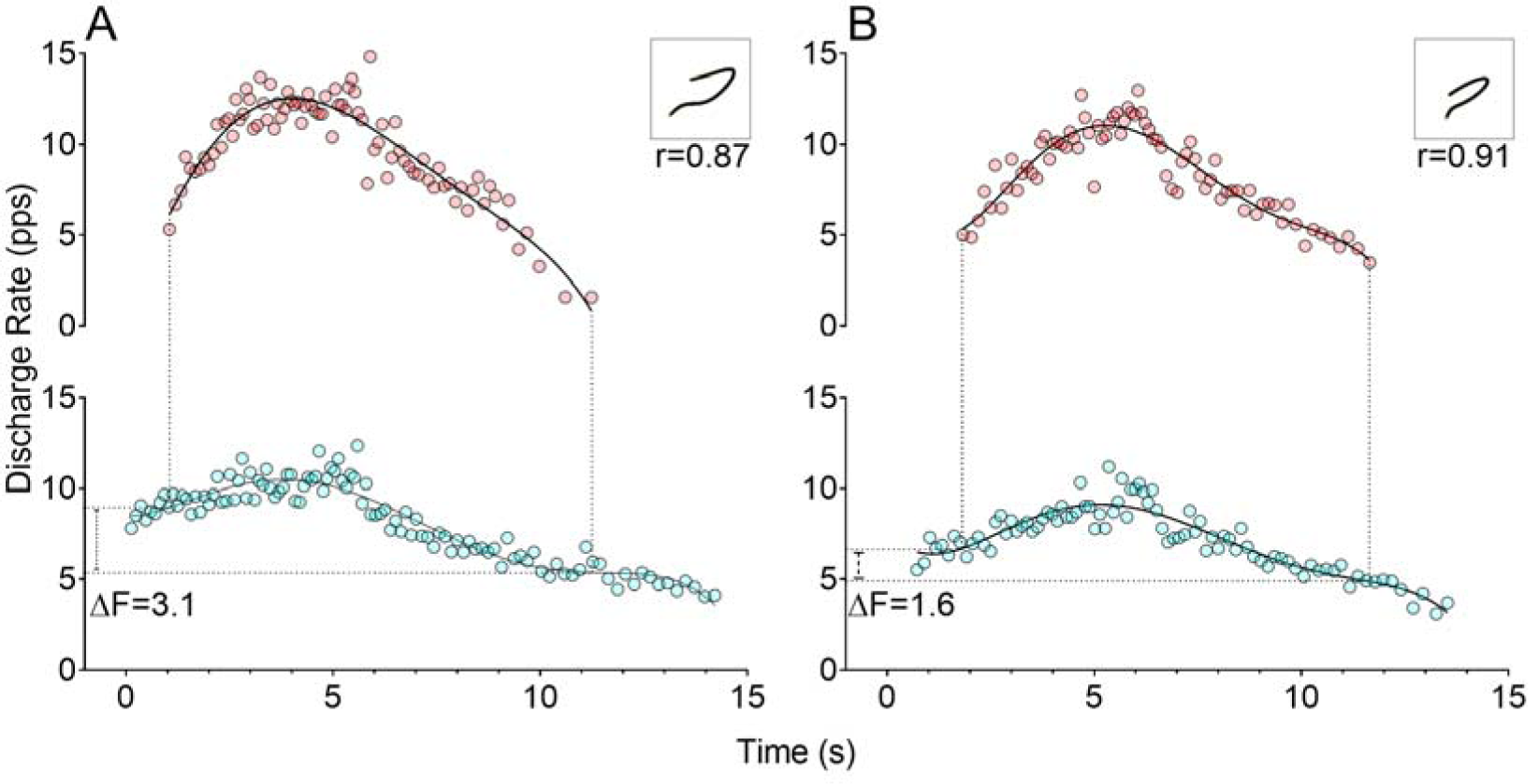
Example of ΔF calculation from soleus motor units before (A) and immediately after passive stretching (B). ΔF was calculated as the difference in discharge rate of the lower-threshold motor unit (blue circles, control unit) at the time of recruitment and de-recruitment of a higher-threshold unit (red circles, test unit). Top right panels (insets) show a strong correlation between the discharge rates of the two units.

### Muscle Stretching Protocol

The stretching procedures were performed on an isokinetic dynamometer with the muscles relaxed. The plantar flexors were stretched five times, with 15-s rest intervals, by rotating the ankle into dorsiflexion at 2°/s until maximal tolerable stretch was attained and then holding at the stretched position for 1 min. This 5-min stretch protocol is the same as used in previous studies reporting reductions in maximal voluntary torque, neural drive to the muscle, and tendon vibration reflexes (Trajano et al. 2014).

### Statistical Analysis

Separate one-way repeated-measures ANOVAs were used to compare changes in ΔFs and peak discharge rates over time (Control 1, Control 2, and Post-stretch). Tukey post-hoc comparisons were performed as follow-up tests. Comparisons between Control 1 and Control 2 were used to investigate changes in the control condition while comparisons between Control 2 and Post-stretch were used to determine changes after the stretching condition. Test-retest reliability (Control 1 and Control 2) of ΔFs and peak discharge rates were evaluated by intraclass correlation coefficients (ICC) via linear regression (Hopkins 2017). Statistical significance was set at an α-level of 0.05. All data are presented as mean and 95% confidence interval.

## RESULTS

### Motor units

The total number of motor units tracked across the three time points was 82 for SOL and 153 for GM. On average 5.5 (3.9 to 7.1) SOL and 10.2 (7.5 to 12.9) GM motor units were tracked per participant. A total of 73 pairs for SOL and 328 for GM motor units were identified across the three time points and used to compute ΔF values. On average 4.9 (1.3 to 8.4) SOL and 21.9 (9.5 to 34.2) GM pairs of motor units were used per participant.

### ICCs

ICC values between Control 1 and Control 2 were high for SOL (0.94, 0.82 to 0.98) and GM (0.97, 0.90 to 0.99) ΔFs. Similarly, high ICC values were observed for SOL (0.95, 0.87 to 0.98) and GM (0.93, 0.79 to 0.97) peak discharge rates.

### ΔFs

There was a significant reduction in SOL ΔFs (F_2,28_=7.63; p=0.002) over time (Control 1 = 2.91 pps (2.02 to 3.79); Control 2 = 2.97 pps (2.03 to 3.92); Post-stretch = 2.23 pps (1.27 to 3.18)), with ΔFs significantly reduced at Post-stretch compared to Control 2 (−25.6% (−42.1 to -9.1); p=0.004; Figure 1). No changes were detected in GM ΔFs (F_2,28_=1.16; p=0.33) over time (Control 1 = 2.89 pps (2.19 to 3.60); Control 2 = 2.86 pps (2.18 to 3.54); and Post-stretch 2.62 pps (1.79 to 3.44)) (Figure 2).

**Figure 2.**
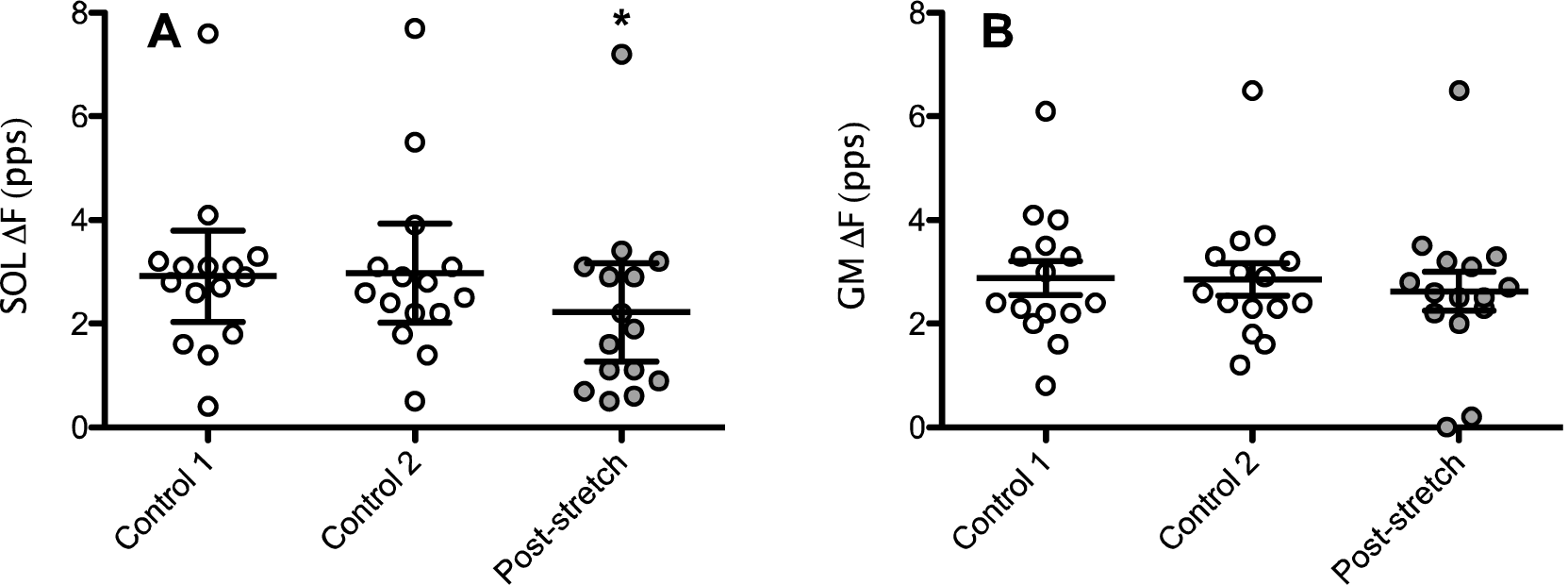
ΔF values of individual participants (n=15) measured in soleus (SOL; panel A) and gastrocnemius medialis (GM; panel B) during the control condition (open circles, Control 1 and Control 2) and immediately after 5 min of passive stretching (gray circles, Post-stretching). Data are also presented as mean and 95% confidence interval. * significantly reduced compared to Control 2 (p<0.01).

### Peak discharge rates

There was a significant increase in peak discharge rates in SOL (F_2,28_ = 5.14; p=0.013) over time (Control 1 = 9.94 pps (9.13 to 10.75); Control 2 = 10.38 pps (9.49 to 11.27); Post-stretch = 10.49 pps (9.61 to 11.38)). However, while peak discharge rates were higher at Post-stretch than in Control 1 (p=0.014), a statistical change was not observed between Control 2 and Post-stretch (p=0.79). No changes were detected in GM (F_2,28_=2.01, p=0.15) motor units over time (Figure 3).

**Figure 3.**
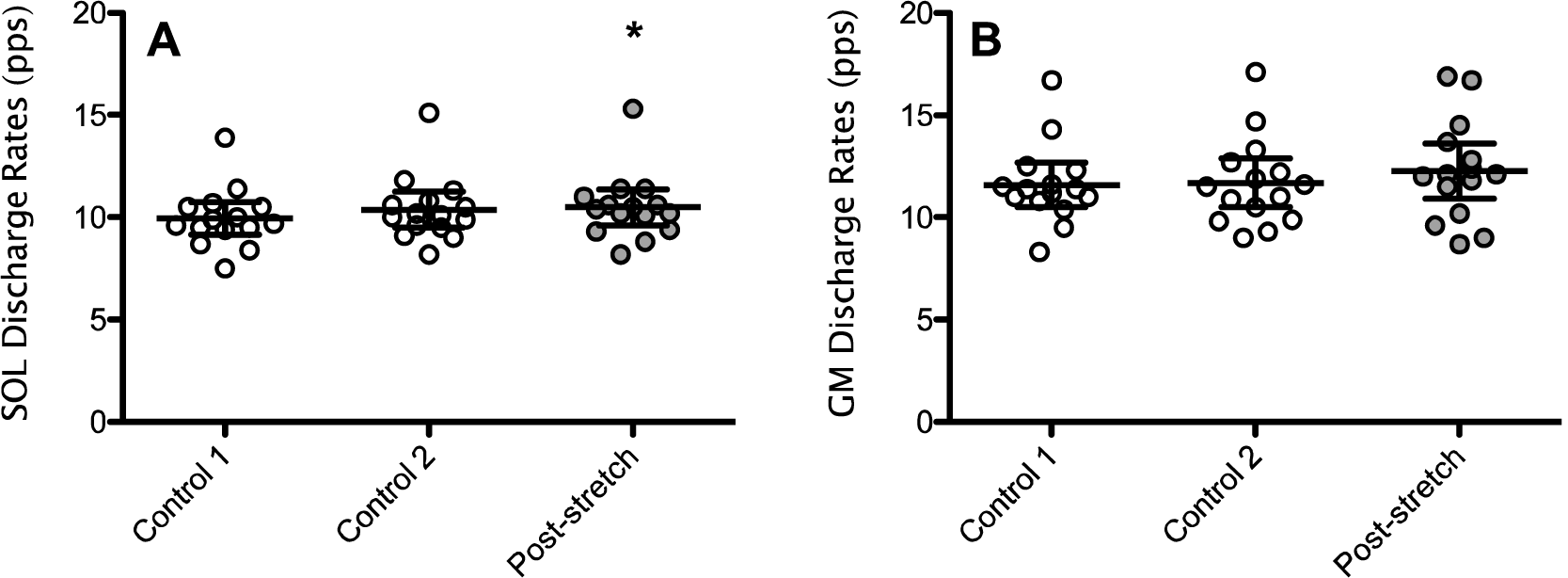
Peak discharge rates for individual participants (n=15) measured in soleus (SOL; panel A) and gastrocnemius medialis; panel B) during the control condition (open circles, Control 1 and Control 2) and after 5 min of passive stretching (gray circles, Post-stretching). Data are also presented as mean and 95% confidence interval. * significantly increased compared to Control 1 (p<0.05)

## DISCUSSION

The main finding of the present study was that passive stretching of the plantar flexors reduced ΔFs in SOL but not GM, suggesting that stretch-induced PIC inhibition occurred only in SOL motor units. These data are not only consistent with the hypothesis that a stretch-induced decrease in PIC strength might underpin post-stretch force reductions.

Motoneuronal disfacilitation has been suggested as a mechanism involved in the reduction in neural drive after passive muscle stretching (Trajano, Nosaka and Blazevich 2017). Preliminary evidence for this hypothesis was gathered in an experiment in which passive stretching reduced the strength of ongoing muscle contractions (i.e. self-sustained motor unit firing) that were elicited via Ia afferent input during tendon vibration and amplified by simultaneous muscle electrical stimulation but which persisted after both tendon vibration and electrical stimulation ceased (28). These data are indicative of a reduction in motoneuron gain, possibly through reductions in PIC amplitudes (Trajano et al. 2014). The present study used a more robust and validated method (the paired motor unit technique) to test this hypothesis. We observed a ∼26% reduction in ΔFs in SOL but not in GM (see Figure 2), suggesting the change caused by passive stretching was very significant, but specific to SOL. In the previous study using tendon vibration it was speculated that inhibition of Ia afferents might be a factor influencing the loss of motoneuron facilitation, as these afferents were the main source of synaptic input in that study (Trajano et al. 2014). However, PIC amplitudes were estimated during voluntary contractions in the present study, where corticospinal projections should be the main source of synaptic input. Therefore, PIC amplitudes appear to be reduced regardless of whether they are estimated as the response to tendon vibration (predominantly Ia input) or using the paired motor unit technique (voluntary contraction).

It is possible that SOL ΔF reduction has a non-localized, ubiquitous, origin. Considerable evidence for this comes from experiments demonstrating that passive stretching of the ipsilateral limb can reduce the motor output of a non-stretched contralateral limb (Caldwell et al. 2019; da Silva et al. 2015; Cè et al. 2020). With respect to a non-localized mechanism, it is interesting to note that passive muscle stretching affects autonomic regulation, changing the sympathetic-parasympathetic balance. More specifically, an increase in parasympathetic and/or decrease in sympathetic drive may be triggered by passive stretching, leading to a post-stretch reduction in noradrenergic activity (Kruse and Scheuermann 2017; Mueck-Weymann, Janshoff and Mueck 2004; Farinatti et al. 2011; Inami et al. 2014). Importantly, PICs are strongly facilitated in the presence of both serotonin and noradrenaline, and PIC amplitude (and therefore ΔF) has been shown to be directly proportional to the level of monoaminergic input from the brainstem (Johnson et al. 2017). For instance, Udina et al. (Udina et al. 2010) found that amphetamine ingestion that increased the presynaptic release of noradrenaline in humans led to a 62% increase in ΔF without changing motor unit initial or average discharge rates (changes in ΔF resulted predominately from changes in decruitment rates). These results are consistent with the results of the present study, in which changes in ΔFs were also observed without changes in peak discharge rates (no difference between Control 2 and Post-stretch; Figure 3). The clear effect of amphetamine ingestion on ΔFs suggests that, similar to what has been observed in animal preparations (Rank et al. 2007; Lee and Heckman 1999), the activation of α1 adrenergic receptors in humans strongly contributes to PIC amplitudes. Therefore, reductions in noradrenergic input from the locus coeruleus after passive stretching could speculatively play a role in the reduction in PIC amplitude. However, more targeted mechanistic experiments are necessary to explicitly test this hypothesis.

The specific reduction that was observed in SOL, but not GM, ΔFs cannot be explained only by the noradrenergic hypothesis and requires further explanation. Several minutes of plantar flexor stretching typically decreases maximal muscle excitation capacity, as measured by EMG amplitude (Trajano, Nosaka and Blazevich 2017). A number of studies have reported this reduction to occur specifically in SOL but not the medial or lateral gastrocnemii, suggesting a potential muscle-specific effect of the stretching (Pulverenti et al. 2020; Pulverenti et al. 2019; Trajano, Seitz, et al. 2013). The reasons for this muscle-specific effect are still unknown but some possibilities deserve consideration. First, PICs tend to be more pronounced in slow-type motor units (Heckman et al. 2008), which are abundant in human SOL (∼80-70%) but less abundant in GM (∼55-60%) (Gollnick et al. 1974; Houmard et al. 1998; Harridge et al. 1996), and have been suggested to play an important role in the tonic firing of postural muscles (e.g. soleus) by decreasing the amount of descending drive required to maintain sustained contractions (Heckman et al. 2008). However, the findings of the present study do not lend support to this assertion as ΔFs were similar between muscles, and lower-threshold motor units are likely to have been sampled from both muscles due to the low intensity of contractions used (20% MVC). Second, muscle spindle Ia-afferent input, which favors slow-twitch motor units and is an important source of PIC initiation, is greater in SOL compared to GM (Tucker and Türker 2004; Eccles, Eccles and Lundberg 1957) and could be negatively affected by stretching (Trajano, Nosaka and Blazevich 2017). However, a problem with this hypothesis is that a lack of change in Ia-afferent pathway excitability is commonly shown after passive stretching (Opplert et al. 2020; Budini et al. 2018; Pulverenti et al. 2020). In particular, one recent study has reported reductions in SOL EMG amplitude without detectible changes in the Ia-afferent pathway (no reduction in H-reflex amplitude) after passive stretching, suggesting that reductions in this pathway are unlikely to contribute to reduction in SOL excitation (Pulverenti et al. 2020). Third, it could be speculated that SOL motoneurons might contain a greater number of PIC-amplifying monoaminergic receptors or a greater density of PIC-producing L-type voltage-gated calcium channels (Ca_V_1.2 and Ca_V_1.3) and/or voltage-gated sodium channels (Na_V_1.1 and Na_V_1.6) (Wilson et al. 2015; Binder, Powers and Heckman 2020). However, little is currently known about these possible differences. Further research is required to determine how PIC strength might be differentially modulated between muscles within a single synergistic group.

In conclusion, passive muscle stretching detectibly reduced PIC strength in SOL but not GM motor units, not only suggesting that a loss of PIC strength might contribute to the loss of muscle force production after passive (static) muscle stretching but that the muscle-specific decrease in muscle activity (EMG) observed previously (Pulverenti et al. 2019) might be explained by a specificity of PIC inhibition. Thus, the present data support the supposition that prolonged periods (e.g. several minutes) of muscle stretching can acutely affect the operation of persistent inward currents in spinal motoneurons, and that this reduces motoneuron (and thus muscle) activation *in vivo* in humans. The development of methods that allow the testing of associations between post-stretch alterations in PIC strength and changes in motor unit firing characteristics during maximal contraction are needed to more explicitly test this hypothesis.

